# Convolutional-LSTM Approach for Temporal Catch Hotspots (CATCH): An AI-Driven Model for Spatiotemporal Forecasting of Fisheries Catch Probability Densities

**DOI:** 10.1101/2024.12.08.627368

**Authors:** Altair Agmata, Svanur Guðmundsson

## Abstract

Efficient fisheries management is crucial for sustaining both marine ecosystems and the economies that heavily depend on them, such as Iceland. Current fishing practices involve decisions informed by a combination of personal experience, current data on environmental and oceanographic conditions, reports from other captains, and target species within the constraints of the fishing quota. However, the intricate spatiotemporal dynamics of fish behaviour make it difficult to predict fish stock distributions. Despite technological breakthroughs in fishing vessel data collection, much of the decision-making still relies heavily on subjective judgment, highlighting the need for more robust, data-driven predictive methods. This paper presents CATCH, a convolutional long short-term memory neural network model that forecasts fish stock probability densities over time and space in Icelandic waters. The framework represents the first utilization of large-scale Icelandic fishing fleet data integrating multidimensional inputs like depth, bottom temperature, and catch data to produce accurate, multivariate forecasts. The model demonstrates high accuracy, low error metrics, and strong structural similarity to observed data, generalizing well across key species such as Atlantic cod, haddock, saithe, golden redfish, and Greenland halibut. Its promising results suggest deep learning models have the potential to optimize fisheries operations, enhance sustainability, and support data-driven decision-making.

## 1 Introduction

Efficient fishing practices are essential for maximizing the economic benefits derived from marine resources while environmental sustainability is safeguarded. This ensures the long-term preservation of both the ecosystem and economic livelihoods of nations heavily reliant on the fishing industry. Iceland, in particular, is a country where the fishing sector plays a pivotal role in its economy and is widely believed to be the country’s single most important industry [1]. In fact, current statistics estimate that the industry contributes approximately 38% of the nation’s total export revenues in 2023, with the total export from fisheries valued at 359 billion ISK in 2023 [2, 3]. The economic significance of the fisheries industry thus underscores the importance of accurate fish stock management strategies. Icelandic waters, influenced by the dynamic North Atlantic currents, are home to commercially significant species, including *Gadus morhua* (Atlantic cod), *Melanogrammus aeglefinus* (haddock), *Reinhardtius hippoglossoides* (Greenland halibut), *Sebastes marinus* (golden redfish), and *Pollachius virens* (saithe). These species, among others, are essential to the economy, with Atlantic cod being one of the most valuable due to its high demand globally and its indispensable role in the ecosystem [4, 5, 2].

In line with the significant role that fisheries play in the economy, the Icelandic fishing industry has evolved to become one of the most technologically advanced of its kind in the world. Over the years, technological innovations have enabled companies to optimize their operations significantly throughout the value chain. These advancements facilitated the collection of massive and diverse datasets that includes information on fishing time, location, sea temperature, depth, and even on catch volume and composition. Currently, skippers gain insights about where and when to fish by manually reviewing and sharing their ships’ data, and combining it with their personal experiences, environmental and oceanographic conditions, and weather forecasts. However, inherent uncertainties in the fishing process limit skippers’ predictive capabilities, which rely heavily on subjective judgment. As a result, the there is a pressing need for an accurate prediction method that guide fishermen toward most efficient and environmentally sustainable fishing at all times [6]. While the current data collected is abundant and rich, existing methods fall short of leveraging its full potential because they lack the necessary analytical capabilities to extract comprehensive predictive insights.

To address these challenges, artificial intelligence (AI) offers promising new technologies, particularly deep learning models, for optimizing operations of fishing fleets [7, 8, 9]. Neural networks, the foundation of deep learning, are models inspired by the biological brain’s structure and can uncover complex, non-linear relationships within the data. They are composed of simple, non-linear modules (neurons) that perform level-wise transformations of increasing abstraction, allowing very complex functions to be learned given enough combinations of such transformations [10]. It is in recent times that AI became the more appealing choice compared to conventional statistical and classical machine learning methods because it can automatically learn non-linear representations of complex-structured data without relying on hand-crafted features or prior domain knowledge [10, 11]. In contrast, conventional statistical and machine-learning techniques were limited in their ability to process natural data in their raw form, requiring rigorous feature engineering along with considerable domain expertise. This advantage has driven the rise of AI applications like ChatGPT in the natural language processing domain, where the model learns directly from raw training data without needing rules in language such as grammar and syntax to be explicitly specified [12, 13, 14, 15].

The spatiotemporal nature of fisheries data requires an architecture that handles both the patterns through time in various temporal scales and spatial relationships across multiple geographical areas. For each of these two tasks, long short-term memory networks (LSTM) and convolutional neural networks (CNN), respectively, are particularly well-suited. LSTM networks, a special type of recurrent neural networks (RNN), are designed to retain information over long sequences of data and thus highly effective for time-series forecasting use cases. The term “long short-term memory” refers to the short-term memory lasting thousands of timesteps that the LSTM architecture aims to provide for RNNs, giving it the ability to analyze long-term sequence dependencies [16]. It also addresses the common vanishing/exploding gradient problem common with RNN’s by keeping track of a separate “cell state” that changes with each iteration [17, 18]. Due to these advantages, LSTM networks are one of the most popular models of choice for sequence-type datasets and have been successfully applied in multiple domains such as finance [19, 20, 21], manufacturing [22, 23, 24], healthcare [25, 26], meteorology [27, 28, 29, 30], fisheries [7, 8, 9]. Meanwhile, CNN’s excel at capturing spatial relationships within the data, making them ideal for analyzing geographical distributions of fish populations. As the name implies, most of the heavy lifting in the feature extraction process within a CNN comes from the convolution operation that happens within each convolutional layer. Within this layer, local conjunctions of features from preceding layers are detected through discrete convolution between learnable kernel filters and the input/intermediate data resulting to the extraction of invariant features such as edges, motifs, parts and objects in an image [10]. These features are then used downstream for classification, typically through a shallow or deep dense neural network. The architecture has seen great success in numerous applications where images are the main input data format [31, 32, 33, 34] or when data can loosely be transformed into image-like representation [35, 36, 37, 22].

An AI architecture that combines the strengths of both CNN’s and LSTM’s thus makes the most sense when tackling the complexity of spatiotemporal fisheries data, particularly when applied to forecasting. Incidentally, such an architecture that literally combines both such frameworks, called Convolutional LSTM, exists and was first applied to precipitation forecasting applications Shi et al. [38]. It has also become an increasingly popular choice and proven effective in applications involving data with both temporal sequences and spatially organized two-dimensional arrays across a variety of fields Sønderby et al. [39], Luo et al. [40], Zhang et al. [41], Behera et al. [42].

This study introduces CATCH (Convolutional-LSTM Approach for Temporal Catch Hotspots) which is an AI model with the goal of producing accurate and timely predictions of fish stock probability densities across space and time in the Icelandic waters. This paper is the first in literature to discuss the use of a deep learning model such as convolutional-LSTM to forecast spatiotemporal fisheries dynamics. This is also the first time that the previously unexplored data collected directly from the Icelandic fishing fleet will be utilized at such a computing scale. Notably, the core objective of this study is to validate the predictive capability of the framework, irrespective of the ecological or fisheries-specific context. This focus is essential because the catch data does not directly capture the biological characteristics of the fish, and external biases such as fishing behavior remain intertwined within the dataset. In Icelandic waters, where environmental variability and fishing pressure can shift rapidly, these AI-driven models open paths for the development of powerful tools for skippers and fisheries operators, to provide them valuable insights for their operations. Our model integrates multidimensional spatiotemporal data—including temperature, depth, and catch per unit effort (CPUE) probability densities—to generate outputs that help optimize fisheries operations, potentially enhancing profitability and long-term sustainability of Iceland’s fishing industry.

## 2 Materials and Methods

### 2.1 Training dataset

The raw data used in this study was a proprietary dataset obtained from various fishing vessels operated by multiple Icelandic fishing companies. These data, encompassing species information, fishing gear, geographical data, vessel information, and environmental parameters, were retrieved using SQL queries to an Amazon Web Services (AWS) Redshift database where it is stored. Due to the proprietary nature of the data, we are unable to disclose the full dataset or make it public.

Before the data was used for model training, the retrieved raw data was first filtered and preprocessed to reduce the complexity of the learning task and make it appropriate to produce the insights and predictions expected of the model. The series of filtering steps taken are enumerated as follows:

1. *Limiting to only bottom trawling (Botnvarpa) as fishing gear type*. Since the main metric used to gain prediction insights on fisheries population is CPUE, this filtering was done to standardize the effort parameter from all observations as different fishing gears have varying catching efficiencies. Bottom trawlers in the dataset had the most amount of observations and also has least efficiency variance within the dataset, thus making it the best choice for effort standardization among different fishing gear types.
2. *Filtering data to 2008 onwards*. The filtering was done to eliminate sampling bias from sparse observations through time as was observed in the data before 2008. Data quality before the cutoff was extremely low and may lead to skewed patterns when used in downstream training.

The filtering process resulted to a dataset with 274,521 spatiotemporal observations spanning across five species: *Gadus morhua* (atlantic cod), *Melanogrammus aeglefinus* (haddock), *Reinhardtius hippoglossoides* (Greenland halibut), *Sebastes marinus* (golden redfish), and *Pollachius virens* (saithe). This dataset is then preprocessed to prepare it for the model training process:

1. *Feature selection*. Features used for training and prediction was limited to three variables: CPUE (kilograms per minute), bottom temperature (celsius), and depth (meters). The temperature and depth were selected, because of their hypothesized impact and data quality. CPUE on the other hand, was obtained by dividing the weight of the catch (kg) with the tow duration (min) for each observation. This normalized the effect of tow duration per catch, thus showing a glimpse of the fish population density at a specific time and area of the catch record. All the ships included in the data have similar average speeds and tow coverage so these factors were not included in the CPUE calculation.
2. *Binning observations across space and time*. The granularity of the observations were reduced through binning and aggregation. Binning was done with respect to space and time, imposing the assumption of local similarity within each bins. In this study, spatial binning was done by 0.5° and 0.25° increments of longitude and latitude, respectively. The binning ranged from -28.75° to -9.25° longitude and 61° to 68.25° latitude, resulting to a 30×40 pixel spatial snapshot for each time observation (Figure 2). Meanwhile, temporal binning was done per month. Binning was set as mentioned to avoid information loss from using excessively large bins and to prevent proliferation of numerous uninformative small bins. Aggregation per bin was done by getting the mean of each group.
3. *Converting features to probability density distributions*. This was done by normalizing each binned spatiotemporal observation for each feature to fractions of unity. Mathematically, it can be expressed as:

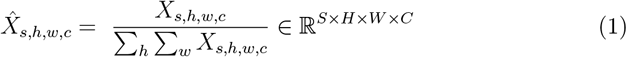

where *X* and 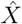 represent the raw and probability density tensors, respectively, of size *S* × *H* × *W* × *C* for each sample observation *s*, latitude *h*, longitude *w*, and feature *c*. Converting the raw feature values to probability densities offers several advantages with respect to forecast modelling, such as enforcing stationarity by normalizing temporal and spatial variability, and focusing on relative distribution dynamics, among others. In the context of CPUE, this normalization means that each value can be interpreted as the fraction of the total catch per unit effort attributed to a specific spatial bin, capturing the relative likelihood of a catch occurring there. This transformation shifts the emphasis from absolute CPUE values to their spatial distribution pattern, thereby mitigating the influence of varying fishing effort and external biases on the predictive model.
4. *Expanding the input tensor to time-lag dimensions*. A common technique used for temporal forecasting predictions to teach the model temporal relationships is by expanding the input tensor to time-lagged version of itself. In this study, time-lag values (*T*) of 3, 6, 12, 16 and 24 months were tested representing various spans of temporal variabilities to be considered per prediction resulting to a final 5D input tensor 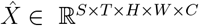.

**Figure 1:**
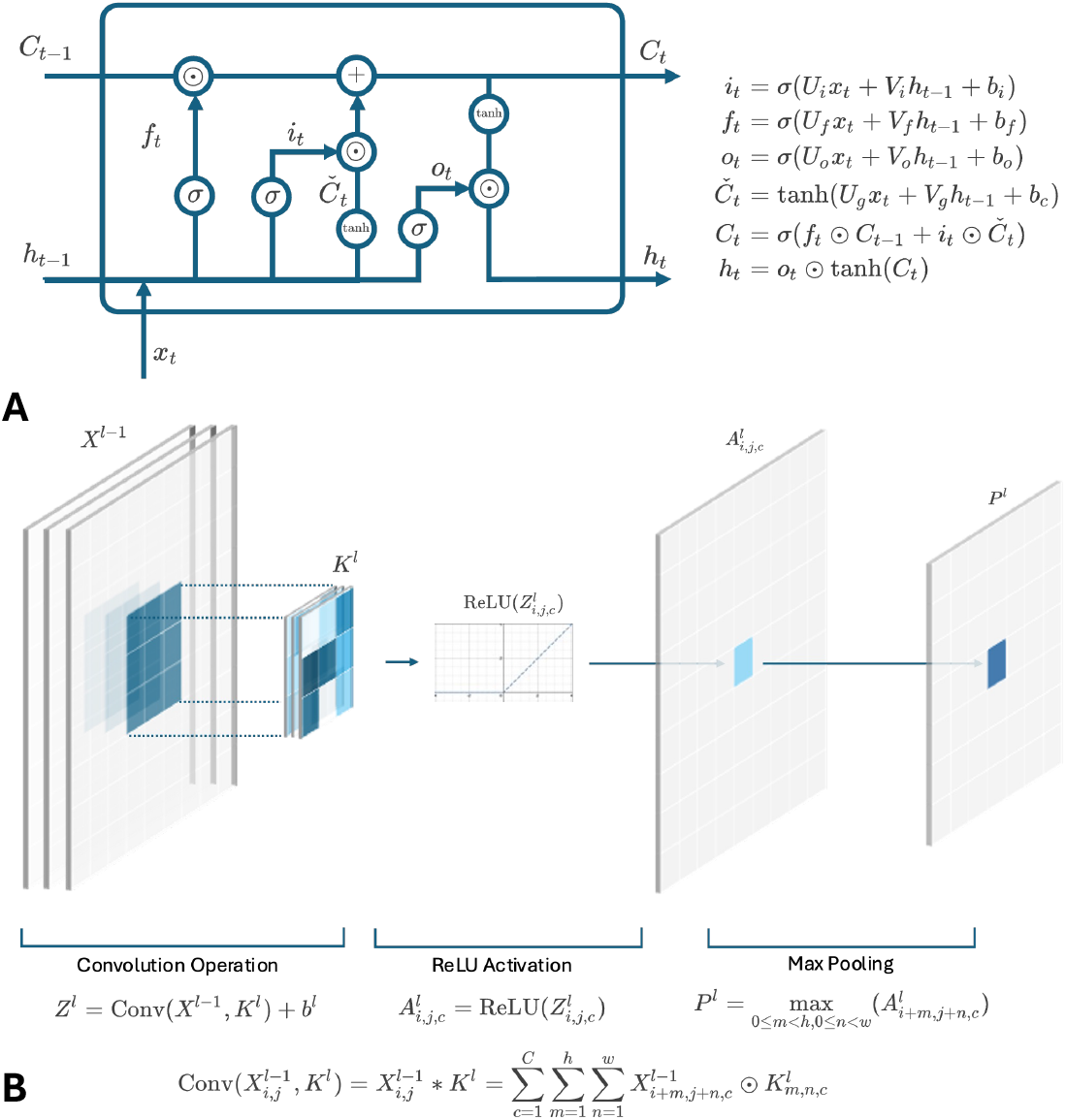
An illustration of the common implementation scheme of A) Long Short-Term Memory (LSTM) neural network [16] and B) Convolutional Neural Network (CNN) [10, 43] along with their corresponding equations.

**Figure 2:**
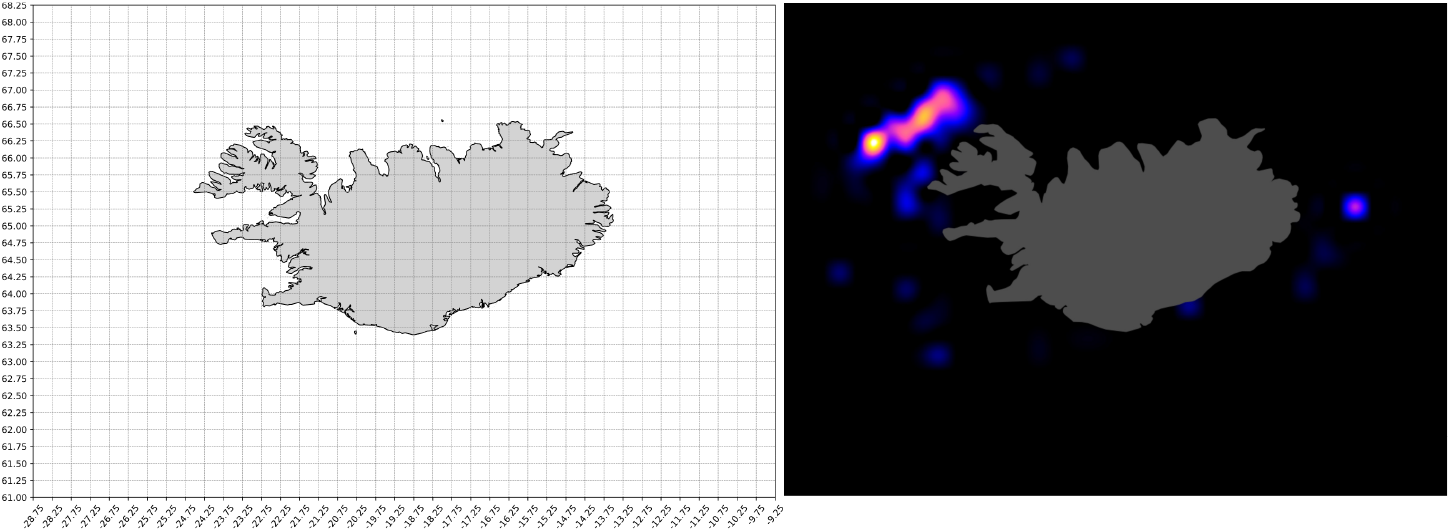
Spatial aggregation of the training dataset showing the binning grid across the Icelandic waters used in the study (left) and the sample snapshot of aggregated CPUE probability density observation with cubic spline interpolation for cod at November 2012 (Right).

**Figure 3:**
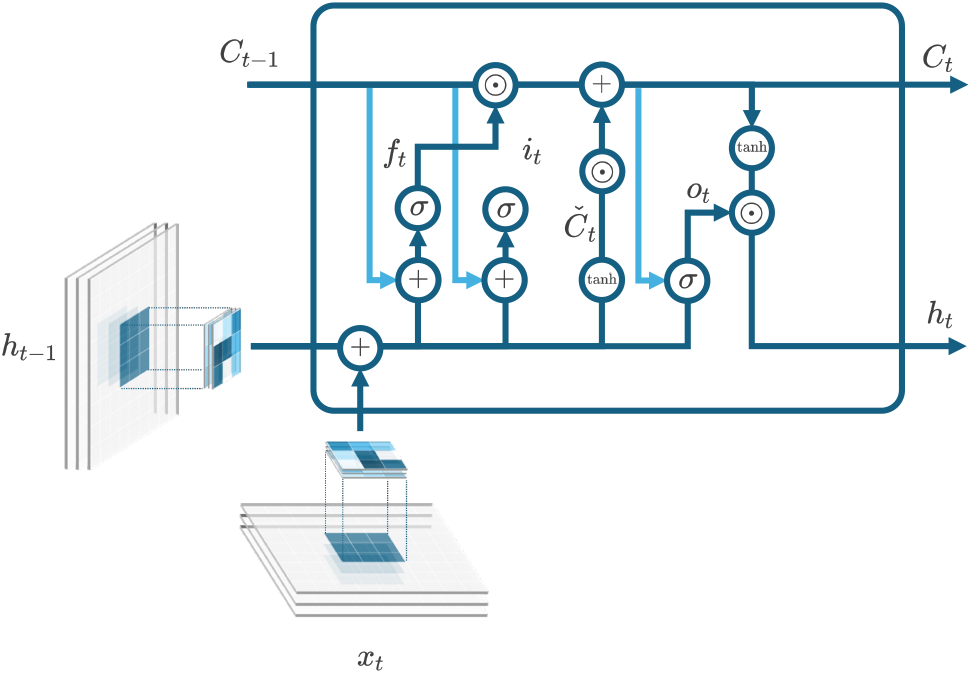
Convolutional LSTM (ConvLSTM) neural network cell used in the study based on [38]

### 2.2 Model architecture

The core component used in the CATCH model was a convolutional LSTM [38] neural network cell designed to capture both spatial and temporal dependencies in fisheries data. The architecture of the model per layer is mathematically expressed as:

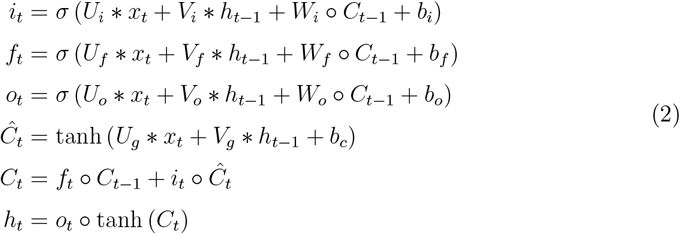

Where ‘∘’ represents the Hadamard product and ‘∗’ the convolutional operation between the trainable kernel weights and an input image described by equation 3 in the case of convolution on input image *x*. In this architecture, the convolution operator is applied in the hidden and input states of LSTM.

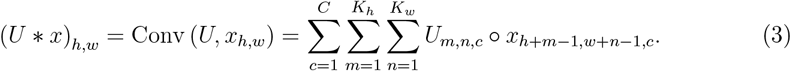

Using the core component as building blocks, a multi-layer model architecture is built representing different layers of data abstraction and expression. Figure 4 shows the implemented architecture for the study wherein each layer’s hidden state is used successively as input towards succeeding ConvLSTM layers. In these layers, expressivity of the data is increased by expanding the feature dimensionality *C* with the use of *F* kernel filters of size *K* with *F > C*. At the last layer, the last hidden state is extracted and passes through the output convolutional layer with *F* 3× 3 kernel filters, compressing the output back to its original dimension *C*. Finally, the convolution output passes through a ReLU activation function to zero out negative probability density predictions and the resulting logit *y*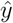 is normalized in a similar way with equation 1:

**Figure 4:**
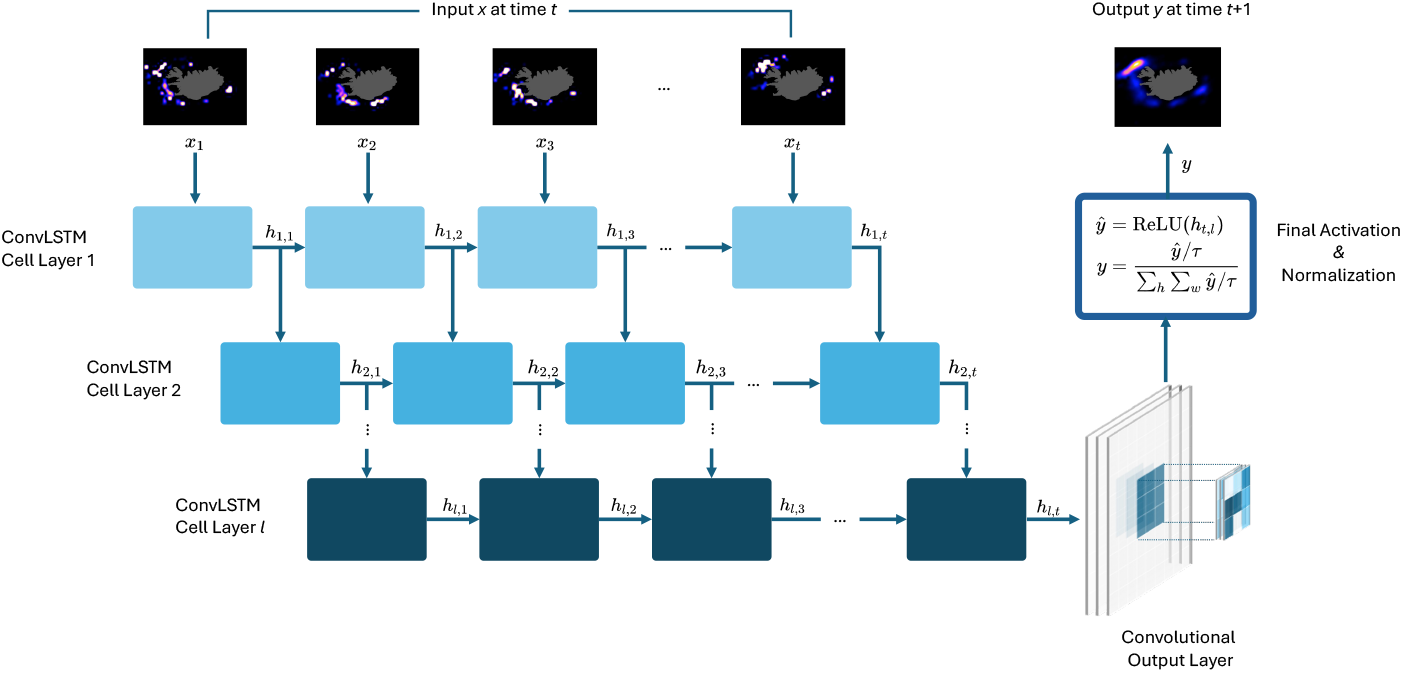
CATCH forecasting model architecture

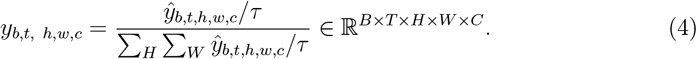

where *τ* is the temperature hyperparameter used to control the entropy of the output probability densities.

### 2.3 Training and hyperparameter tuning

The model is trained through back-propagation through time (BPTT) [44] by minimizing a weighted binary cross-entropy (BCE) loss which is used to prioritize the accuracy towards the CPUE probability density prediction. For each feature *c*, the BCE loss for a single element is given by:

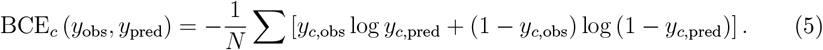

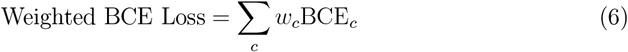

In this study, the weights *w*_*c*_ for each feature used are 0.6, 0.2 and 0.2 corresponding to CPUE probability density, bottom temperature, and depth, respectively.

The model was trained using data of varying lag size, and thus model context window, to gain prediction insights from variations attributed to short-term and long-term patterns. Specifically, context sizes of 3, 6, 12, 18, and 24 months backward were used in the study but only focusing on cod (*G. morhua*) first as this is the species with the most number of observations within the dataset. The context length with the best results are then used to train and predict across all the five species. Each of the dataset was split to train, validation and test set, where the validation and test sets are set as 12-months-worth of data each covering 06-2022 to 05-2023 and 06-2023 to 05-2024, respectively. Hyperparameters for each of the training run were obtained automatically using Bayesian optimization through the Keras tuner library across 100 trials with the objective of minimizing the validation loss. The hyperparameter values/ranges used in the hypertuning process were summarized in Table 1. Decisions about the ranges and fixed values were made based on prior experiments, computational efficiency, and available compute resources.

**Table 1:** Hyperparameter values/ranges explored in the study.

| Hyperparameter | Value / Range |
| --- | --- |
| Number of filters ($F$) | [4, 6] |
| Kernel size ($K$) | $3 \times 3$ |
| Number of ConvLSTM layers ($L$) | [2, 3] |
| Dropout in ConvLSTM layers | [0.1, 0.5] |
| Temperature ($\tau$) | [0.2, 0.7] |
| Max epoch ($E$) | 50 |
| Batch size ($B$) | 32 |
| Learning rate ($R$) | $[1 \times 10^{-5}, 1 \times 10^{-3}]$ |

### 2.4 Model evaluation

Predictions are evaluated based on performance with regard to the model’s one-at-a-time (OAT) and recursive forecast. OAT prediction refers to predicting only one month at a time and that at every timestep, the model uses an input comprised purely of observed instances. In contrast, recursive forecast is done through generation of sequential predictions for a specified forecast length by iteratively using the model’s previous predictions as inputs for subsequent predictions, creating a chain of forecasts over time while maintaining a consistent input size through slicing. Such predictions are evaluated against the testing data using metrics summarized in Table 2.

**Table 2:**
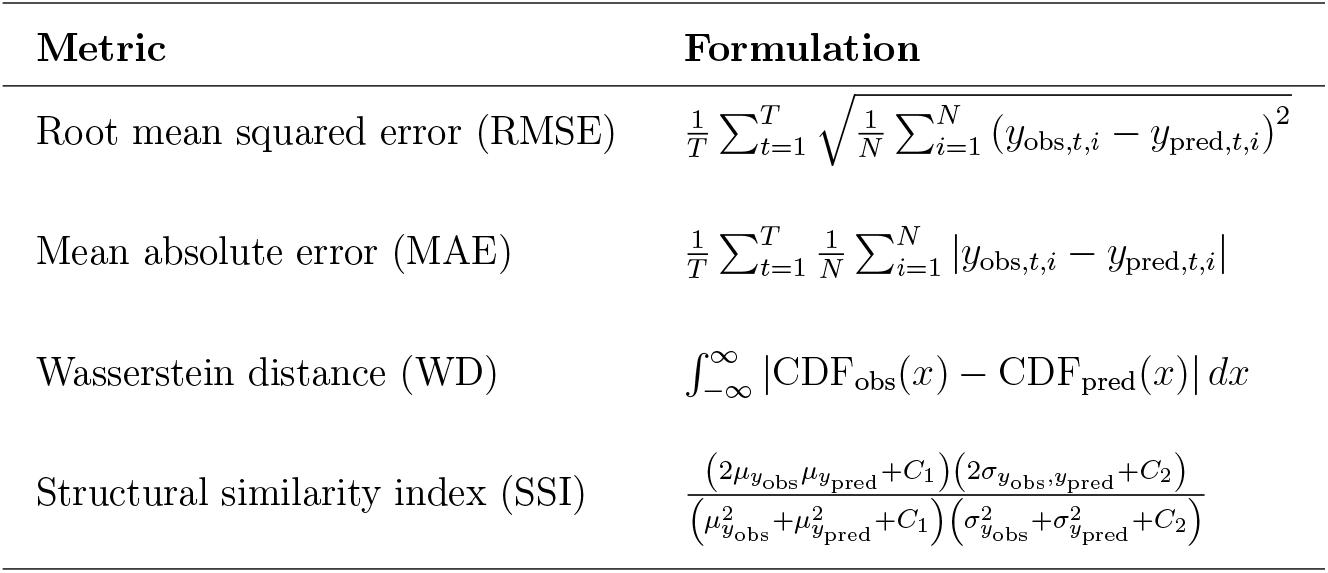
List of metrics used to evaluate model performances and their corresponding mathematical formulations.

RMSE and MAE are standard regression metrics that measure error with varying sensitivity to outliers. Relative fit metrics such as percent error (MAPE) are unsuitable in this context because the response variable frequently contains zero or near-zero values, which can lead to disproportionately large or even infinite percent errors. Although the problem could be treated as a regression task, both the input data and model output are interpreted as probability densities. Consequently, the study employs a suite of evaluation metrics, each offering a distinct mathematical perspective on performance. In particular, the Wasserstein distance (WD) is used to quantify the optimal transport cost required to transform one distribution into another [45] approaching the evaluation as comparison between probability density distributions. Additionally, Structural Similarity Index (SSI) is incorporated, as it is widely used in computer vision to assess differences in intensity, contrast, and structural covariance between images [46], thereby providing a measure of structural consistency across spatiotemporal frames.

## 3 Results

Table 3 and Figure 5 summarizes and illustrates the performance matrix on *G. morhua* test data across varying lag sizes (*L*). A close concurrence is observed between the model’s predictions and the actual observed values as consistently low RMSE and MAE values were observed. Particularly for OAT predictions, the MAE varies from 1.07 × 10^*−*3^ to 1.25 × 10^*−*3^ across the different lag sizes, while the RMSE varies from 4.55 × × 10^*−*3^ to 4.88 × 10^*−*3^. On the other hand, the RMSE and MAE values for recursive predictions range from 4.56 × 10^*−*3^ to 5.16 × 10^*−*3^ and 1.08 × 10^*−*3^ to 1.38 × 10^*−*3^, respectively. Low WD values, ranging from 0.80 × 10^*−*3^ to 1.00 × 10^*−*3^, and high SSI values, ranging from 0.95 to 0.96, were also observed supporting the CATCH models robustness and accuracy. This is despite the striking difference between the nature of the observed and the predicted datasets (Figure 6 and Figure 7).

**Table 3:**
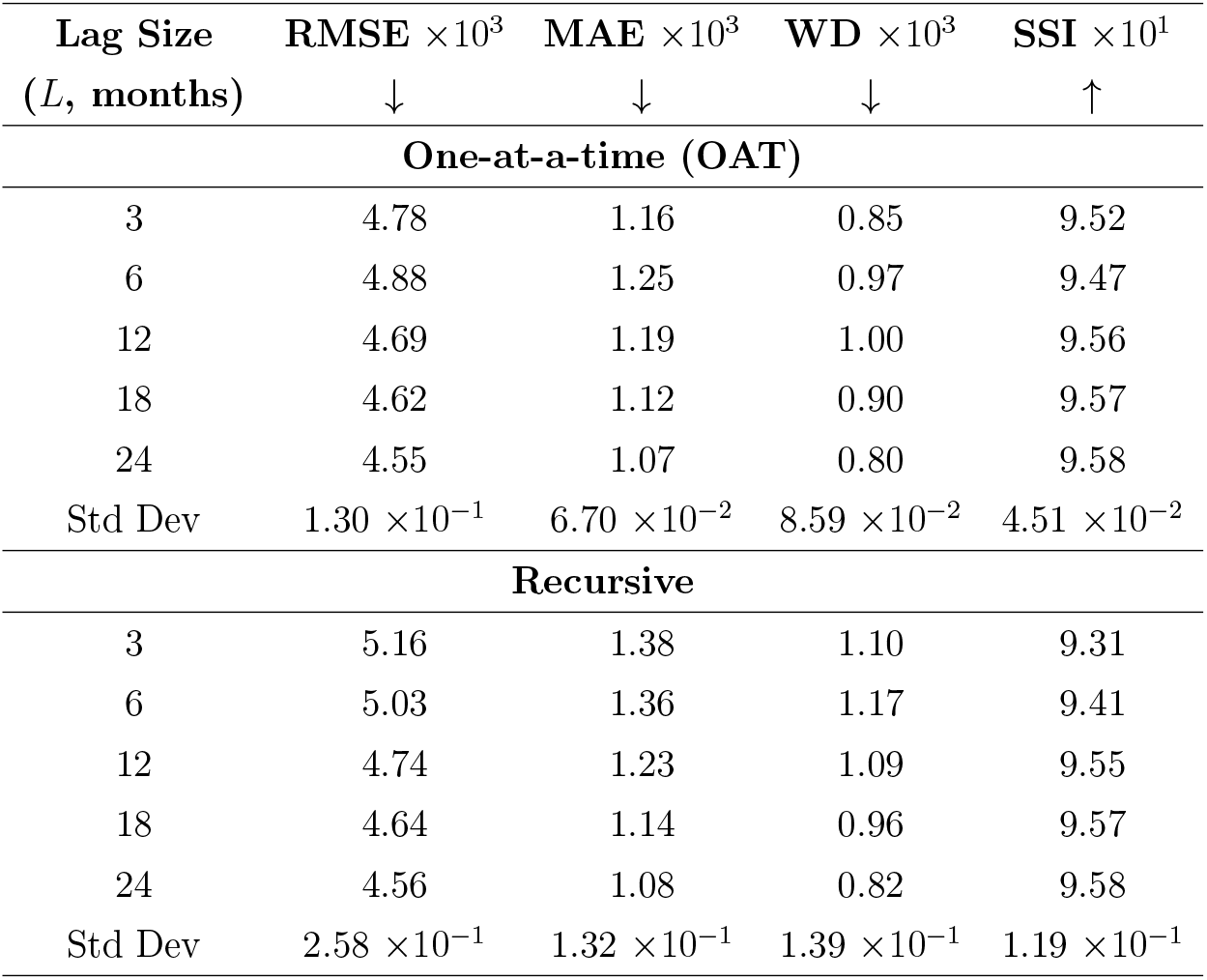
Specific metric values for OAT and recursive predictions for *G. morhua* with respect to observed values as functions of lag size (*L*) from 2023-06 to 2024-05 data. Arrows indicate direction of better scores, i.e. down means the lower the better and up means the higher the better. Standard deviation are reported in absolute unscaled units for easier comparison across varying orders of magnitudes.

| Lag Size<br>($L$, months) | RMSE $\times 10^3$<br>↓ | MAE $\times 10^3$<br>↓ | WD $\times 10^3$<br>↓ | SSI $\times 10^1$<br>↑ |
| --- | --- | --- | --- | --- |
| **One-at-a-time (OAT)** |  |  |  |  |
| 3 | 4.78 | 1.16 | 0.85 | 9.52 |
| 6 | 4.88 | 1.25 | 0.97 | 9.47 |
| 12 | 4.69 | 1.19 | 1.00 | 9.56 |
| 18 | 4.62 | 1.12 | 0.90 | 9.57 |
| 24 | 4.55 | 1.07 | 0.80 | 9.58 |
| Std Dev | $1.30 \times 10^{-1}$ | $6.70 \times 10^{-2}$ | $8.59 \times 10^{-2}$ | $4.51 \times 10^{-2}$ |
| **Recursive** |  |  |  |  |
| 3 | 5.16 | 1.38 | 1.10 | 9.31 |
| 6 | 5.03 | 1.36 | 1.17 | 9.41 |
| 12 | 4.74 | 1.23 | 1.09 | 9.55 |
| 18 | 4.64 | 1.14 | 0.96 | 9.57 |
| 24 | 4.56 | 1.08 | 0.82 | 9.58 |
| Std Dev | $2.58 \times 10^{-1}$ | $1.32 \times 10^{-1}$ | $1.39 \times 10^{-1}$ | $1.19 \times 10^{-1}$ |

**Figure 5:**
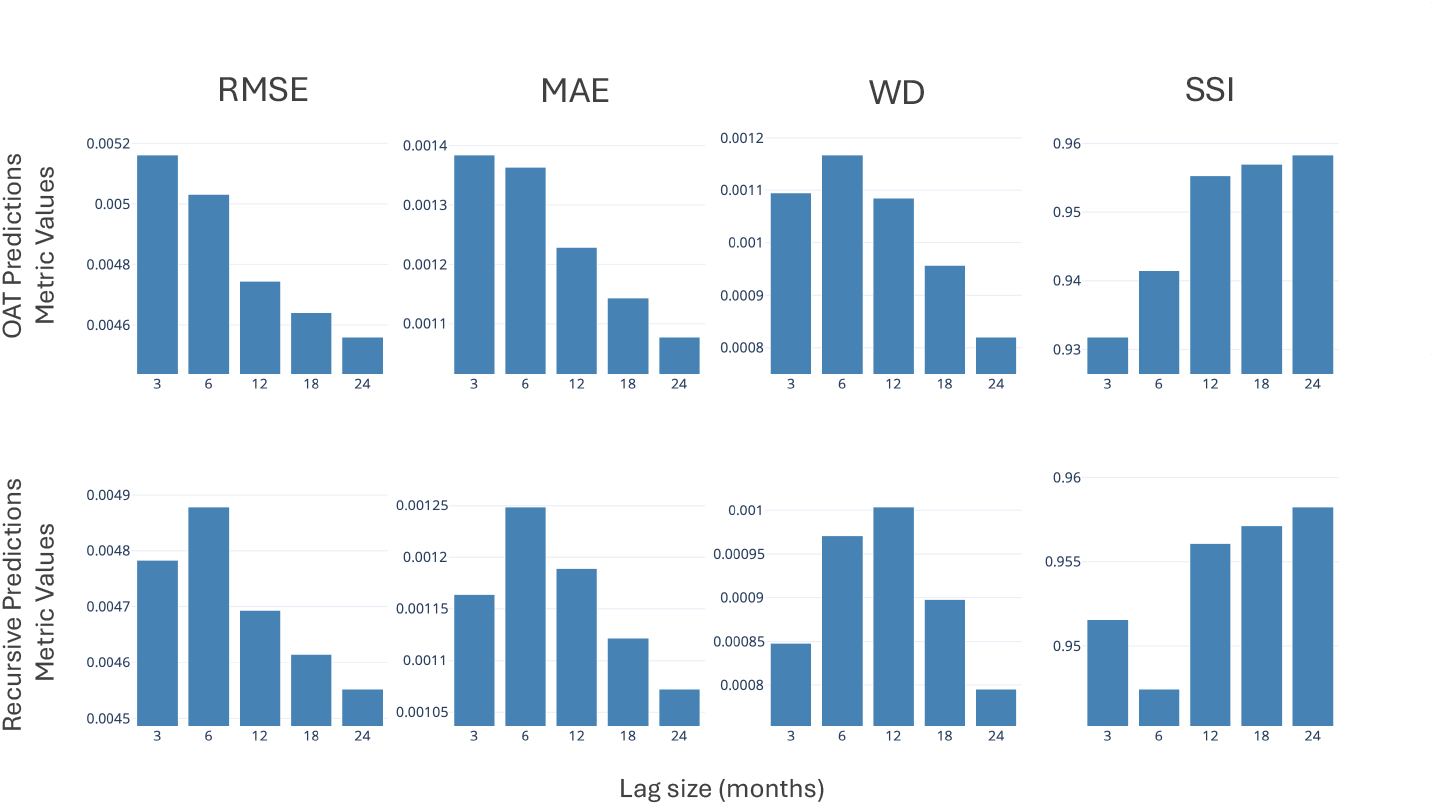
Metric values for one-at-a-time (OAT) and recursive predictions for *G. morhua* with respect to the observed values as functions of lag size (*L*, months) from 2023-06 to 2024-05 data.

**Figure 6:**
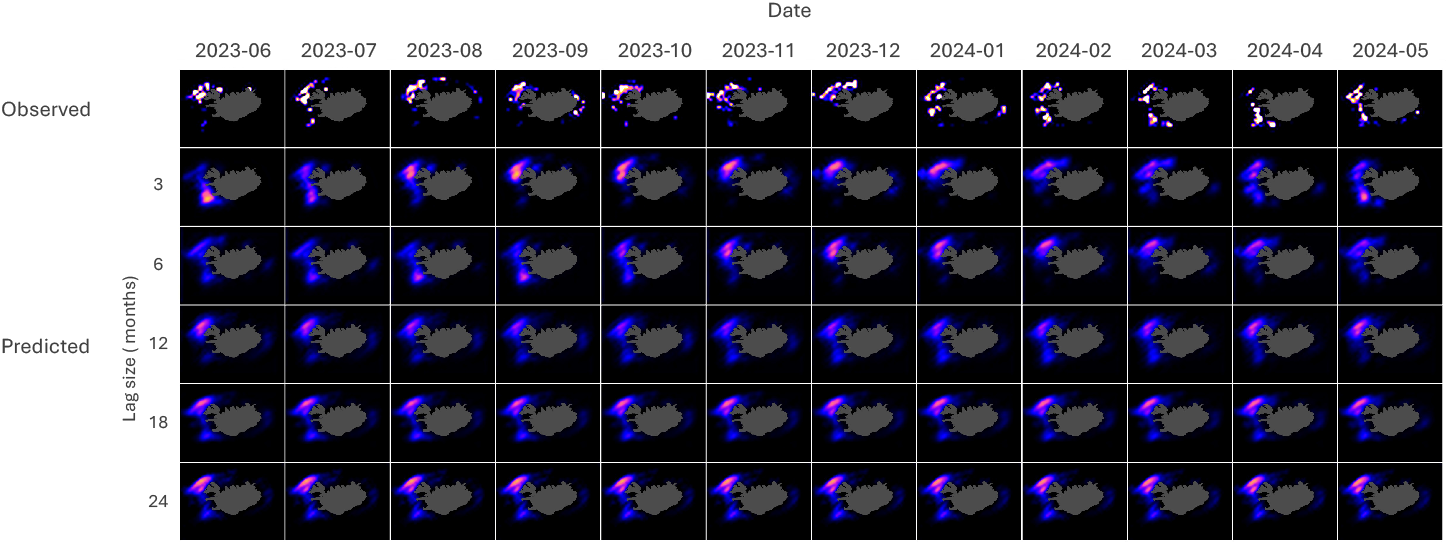
OAT forecast results for *G. morhua* on the test data from 2023-06 to 2024-05 across varying lag sizes. Higher intensities depict higher probability density.

**Figure 7:**
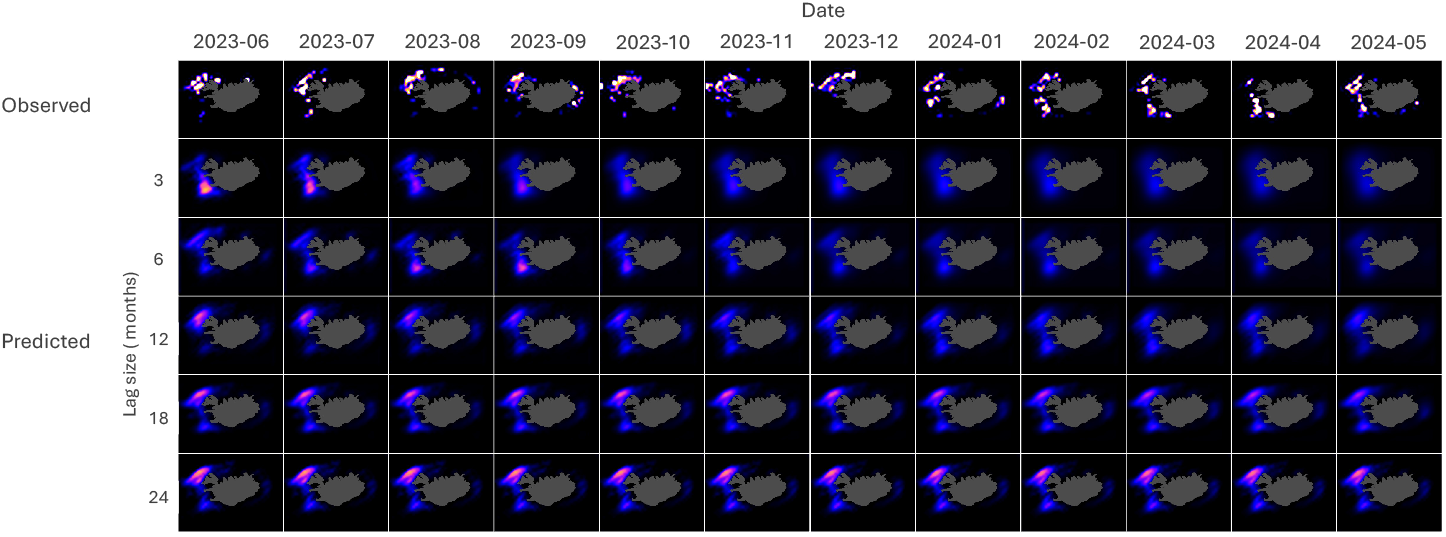
Recursive forecast results for *G. morhua* on the test data from 2023-06 to 2024-05 across varying lag sizes. Higher intensities depict higher probability density.

Figure 6 and Figure 7 shows the plotted observed catch (top row) and the predicted probability catch (bottom rows) in months for both OAT and recursive forecasts, respectively. These figures highlight the difference between the nature of the observed and predicted probability catches—with the observed datasets seen as highly discontinuous with multiple local clusters and the predicted datasets as smooth and continuous. The discontinuous nature of the observed dataset is due to the incomplete and uneven sampling from the fishing fleets. The figures also highlight the movements within the data across space and time and how different *L* values affect these movements in the predictions. While general concordance of the predicted with observed data on probability density distributions is apparent for all the lag sizes, movement is more obvious in models with lower *L* values and slowly diminishes as *L* increases.

Table 4 summarizes performance metrics of test data from *P. virens, S. marinus, R. hippoglossoides*, and *M. aeglefinus* with a lag size of 3 months. Similar patterns of close concurrence are observed between the model’s predictions and the actual observed values as consistently low RMSE and MAE values were observed in order of 10^*−*3^. Low WD values, in the order of 10^*−*3^, and high SSI values, close to unity, were also observed. Figure 8 illustrates the observed and predicted probability densities across time for all the four species. It also highlights how the alignment of predictions with the observed data, showcasing a strong correspondence and accurate reflection of overall spatiotemporal patterns despite varying dynamics across species.

**Table 4:** Specific metric values for OAT with respect to observed values across various species (*P. virens, S. marinus, R. hippoglossoides*, and *M. aeglefinus*) from 2023-06 to 2024-05 data with lag size of 3 months. Arrows indicate direction of better scores, i.e. down means the lower the better and up means the higher the better.

| Species<br>Name | RMSE $\times 10^3$<br>↓ | MAE $\times 10^3$<br>↓ | WD $\times 10^3$<br>↓ | SSI $\times 10^1$<br>↑ |
| --- | --- | --- | --- | --- |
| *P. virens* | 5.69 | 1.31 | 1.11 | 9.41 |
| *S. marinus* | 5.23 | 1.11 | 0.83 | 9.58 |
| *R. hippoglossoides* | 8.11 | 1.24 | 0.96 | 9.70 |
| *M. aeglefinus* | 5.33 | 1.27 | 1.01 | 9.38 |

**Figure 8:**
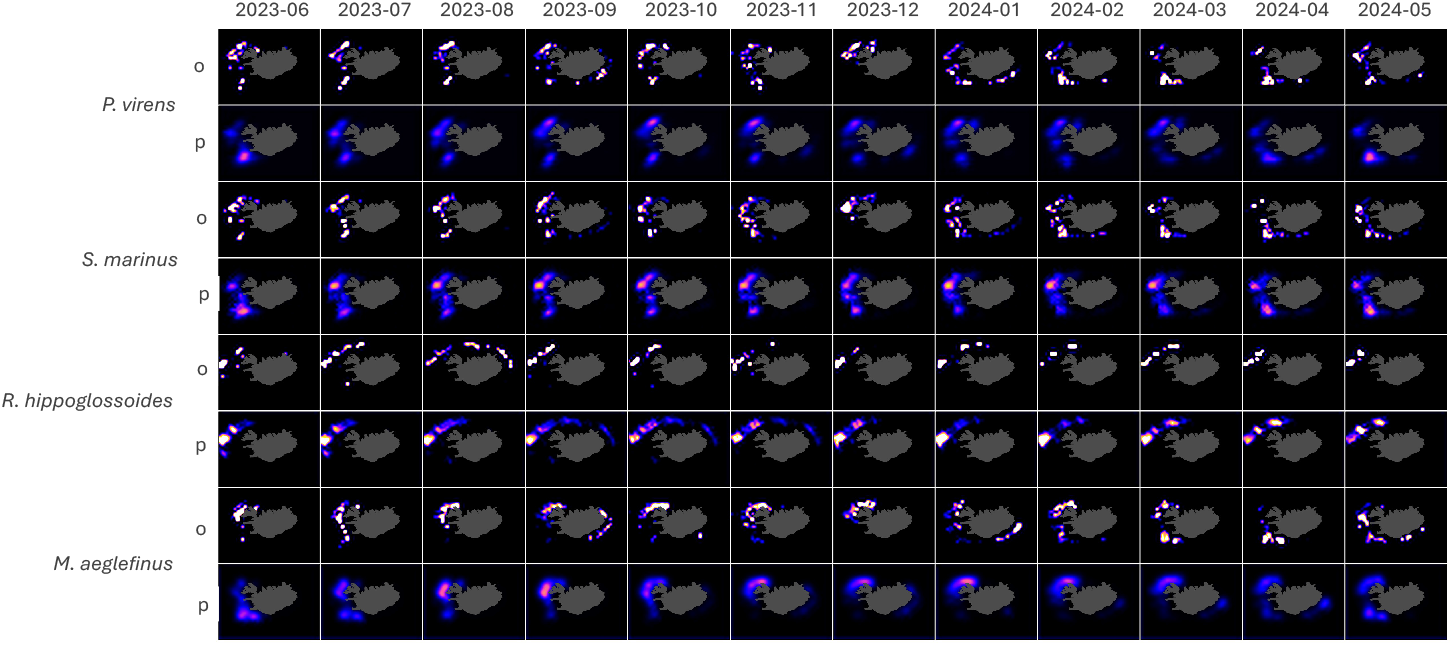
OAT forecast results (*p*) for four species (*P. virens, S. marinus, R. hippoglossoides*, and *M. aeglefinus*) against the test data (*o*) from 2023-06 to 2024-05 with lag size of 3 months. Higher intensities depict higher probability density.

## 4 Discussion

### 4.1 Model performance and evaluation

The results of the experiment imply that the CATCH model does a good job at capturing the data’s underlying spatial and temporal patterns. This is exemplified by the favorable performance metrics for the one-at-a-time (OAT) and recursive forecast predictions as shown in Table 3 and Figure 5. Consistently low RMSE and MAE values, which are in the order of 10^3^, implies that the model’s predictions closely concur with the actual observations.

The CATCH model’s low WD values, in the order of 10^*−*3^, also imply that the predicted probability densities closely match the observed distributions, which is crucial for accurate probabilistic forecasting. Despite the difference in nature of the observed and predicted datasets, low WD values shows that the model does not overfit and generalizes the patterns well, producing predictions that align with ecological intuition rather than overfitting to patches of data. This ensures the model’s capability to extend predictions to unobserved regions, filling data gaps in a plausible manner while ensuring that the predicted distributions maintain structural coherence and environmental relevance. Indeed, this is also supported by the high Structural Similarity Index (SSI) values approaching unity, demonstrating that the model maintains strong structural fidelity in its forecasts across time and signifies that the model preserves important spatial feature structures when compared to the observed data over time. Overall, these results highlight the model’s robustness in capturing both spatial and temporal dynamics while generating predictions that are not only accurate but also ecologically meaningful, making it a valuable tool for potentially understanding and managing fisheries populations.

### 4.2 Comparison of OAT and recursive forecasts

When comparing the metric values between OAT and recursive predictions, OAT forecasts consistently exhibits lower RMSE, MAE, and WD, along with higher SSI. This highlights the nature of compounding errors in recursive forecasting, as the model gradually uses its own synthetic predictions as input data over time. Additionally, examining the metrics for *L* = 24 reveals that both OAT and recursive predictions yield values that are close to each other with minimal differences. This trend suggests that as *L* becomes larger, the recursive metrics converge towards those of the OAT predictions. This thus implies that at least in the context of the training data used in the study, utilizing a context length of at least 24 is recommended in order for recursive forecasts to achieve reliability comparable to OAT predictions. Such observations align with literature findings [47, 48] suggesting that long-range recursive forecasting requires a larger context window.

### 4.3 Effect of lag size

The general trend observed on OAT predictions is that of improving performance as the number of lag size increases. This might indicate that a larger lag size, corresponding to a broader context window and input information, provides the model with wider context per prediction, enabling more accurate long-term forecasts. However, looking at the trend for the WD metric —which considers the evaluation as a comparison between probability densities—it appears that higher model performance might also be obtained through smaller context windows. Indeed, the plot on recursive predictions (Figure 5) also shows a similar pattern, where the performance initially declines with increasing lag size and then starts to improve after *L* = 12. This suggests that two independent temporal patterns are at play, such that increasing or decreasing *L* leads to a trade-off between them. Such is a common scenario in time-series modeling where temporal dependencies are broadly classified into long-term and short-term impacts, typically reflected as trends and fluctuations, respectively [47]. In this case, lower lag values might be more sensitive to short-term dynamics and thus captures them well, despite not reliably capturing longterm relationships. If such a trade-off pattern exists within the dataset used in the study, then the opposite should be true for a model with higher *L*. Results in Figure 6 provide visual affirmation of this reasoning. As *L* increases, spatial movement and intensity variations of the hotspots become less dynamic. In the observed test data, movement of the hotspots through time from north to south are seen to happen briefly from June to July 2023, and then slowly manifests again from January to May 2024. Such temporal dynamics is seen to be captured only by models with *L* of 3, 6 and 12 with movement diminishing as *L* increases. While increasing lag time improves the metrics and can be beneficial for capturing overarching patterns, it may lead to the attenuation of significant short-term events, which are critical in certain forecasting applications.

### 4.4 Multi-species applicability

Building on the emphasis on short-term dynamics and considering that the observed data covers only 12 months—where such dynamics are more apparent—the model with *L* = 3 was chosen to predict across the remaining four species (*P. virens, S. marinus, R. hippoglossoides*, and *M. aeglefinus*). This decision was made because it demonstrated the best performance metrics and visual concordance with respect to capturing the movements within the test data, as evidenced by its performance with *G. morhua*. Figure 8 and Table 4 present the results of the CATCH model predictions for these species. From the figure, it is evident that the model effectively grasps the spatiotemporal movements shown by the observations, similar to the performance seen in *G. morhua*. The movement of hotspots and intensity variations over time are well-represented, indicating the model’s proficiency in modeling essential spatial dynamics for each of the distinct datasets. Quantitatively, this is supported by metric performances in Table 4 where the values for RMSE, MAE, WD and SSI fall within similar orders of magnitude as those for *G. morhua*. These consistent metrics across multiple species demonstrate the robustness of the model in capturing the patterns even across varying training data distributions exhibited by different species.

The robustness of the model’s framework suggests that the model potentially possesses the capability to generalize fish species behavior, at least for bottom-dwelling species that are being represented in this study. Having a single model that can effectively forecast multiple species behavior is particularly significant, as it implies that the model can capture underlying patterns and movements characteristic of the species as a group. Such generalization capability enhances the model’s utility, potentially reducing the need for species-specific models and extensive amounts of training data for each species, as training can be done using only a handful of representative species to generalize patterns within a species group. This would make the model a valuable tool for predicting distributions of under-represented species in data-limited contexts. Moreover, the model’s proficiency in capturing multi-species interactions could have important implications for ecosystem management and conservation efforts. By accurately modelling and subsequent extraction of model explainability insights on multi-species interactions, the CATCH model can provide valuable insights into species distribution patterns, habitat utilization, behavioral correlations, and potential areas of ecological significance. However, it is important to acknowledge potential limitations of the framework used within the study. While the model performs well with the species tested, further validation with additional species and longer observation periods would strengthen confidence in its generalization capabilities. Furthermore, the training and testing data used in the study was not validated in the context of known ecological dynamics from literature sources as the focus was solely the model’s capability to produce accurate forecast given the observations in the data. Future work could thus involve generating and aligning insights with more context on the fish biology of these representative species, testing the model with other species group that exhibits completely different movement patterns, such as pelagic species, and training one general model from crafted features that capture multi-species interactions.

## 5 Conclusion and recommendations

This study introduced the CATCH model, a deep learning framework utilizing Convolutional LSTM networks to forecast fish stock probability densities in Icelandic waters. This is the first in literature to discuss the utilization of this framework to the previously unexplored data collected by the Icelandic fishing fleet for the hope of aiding the Icelandic fishing industry and environmental sustainability. The model demonstrated high accuracy and robustness in predicting the spatial and temporal distributions of key commercial fish species, as evidenced by low RMSE, MAE, and WD, along with high SSI scores. These metrics show the generalization capability of the model without overfitting and how well it captures intricate spatiotemporal patterns. Interestingly, the model demonstrated potential to generalize across several species, accurately representing the distribution patterns of Greenland halibut, Atlantic cod, haddock, saithe, and golden redfish. This implies that the model might be applied to different species groups, which would eliminate the requirement for species-specific models and make it a useful tool for managing fisheries in situations when data is scarce.

Future recommendations for addressing the limitations and gaps identified within the study may include:

1. *Testing Multi-Species Generalization*. Further validation of the model’s generalization capabilities should be conducted by testing it on additional species, including those with different behavioral patterns, such as pelagic species. This will help determine the model’s applicability across diverse species groups. Moreover, the training experiments performed in the current study involved independent CATCH models individually trained from different species and then subsequently used to predict across each of these species. A general model, while more computationally demanding and theoretically more complex, should be trained from a single dataset with crafted features that inherently highlights patterns for multi-species interactions. This could, in turn, forecast distributions even outside the representative species in the training data.
2. *Incorporating Fish Biology Context* Aligning the model’s insights with detailed fish biology (such as fish length and weight distributions) and ecological data can enhance its predictive power and interpretability. Integrating biological factors such as spawning cycles, feeding habits, and migration patterns could provide a more comprehensive understanding of species distributions.
3. *Enhancing Model Inputs*. Exploring additional environmental variables, such as salinity, ocean currents, wind speed, and wind direction, among others, may improve the model’s accuracy and provide deeper insights into the factors influencing fish stock movements.
4. *Long-Term Forecasting* : Extending the model to perform long-term forecasts could be valuable for strategic planning in fisheries management. This would involve training the model with longer time-series data and testing its ability to predict future stock distributions over extended periods.

Enhancing the CATCH model in this way could significantly increase its value as a tool for operational research in the fishing industry by boosting potential profits and increasing the sustainability of fishing operations. Equally important, the model can provide deeper insights into the robustness of individual fish stocks, enabling better strategic measures to ensure sustainability and the preservation of newly spawned generations.

## 6 Acknowledgement

We gratefully acknowledge the invaluable contributions of Dame Loveliness Apaga-Agmata and Elías Elíasson, whose insightful feedback and thorough critique greatly enhanced the quality of this paper. Special thanks to Kristrún Arnarsdóttir for her contributions through insightful discussions, brainstorming, and careful reading of the manuscript. We also extend our gratitude to Guðmundur Kristjánsson and Runólfur V. Guðmundsson for their thoughtful insights, as well as to Arnar Kristjánsson for his practical perspectives and expertise as a skipper, which enriched the study.

We are deeply thankful to the organizations that provided crucial data for this work: Brim hf., Útgerðarfélag Reykjavíkur hf., Samherji hf., Síldarvinnslan hf., and Guðmundur Runólfsson hf. Their support was instrumental in making this research possible.

## 7 Author contributions

A.A. and S.G. were both involved with the conception of ideas, data collection, designing the methodology, and analyzing the results. A.A. executed the modeling aspects, and prepared the manuscript. S.G. reviewed the manuscripts, gave critical feedbacks, and ultimately approved the paper for publication.

## 8 Conflict of interest

The authors declare no conflict of interest.

## 9 Funding

This study was supported by funding from Brim hf. and Útgerðarfélag Reykjavíkur hf., whose contributions were instrumental in facilitating the research and enabling access to proprietary data essential for the project. We gratefully acknowledge their generous support and commitment to advancing this work.

## 10 Data availability

The data underlying this study are proprietary and were provided by Brim hf., Útgerðarfélag Reykjavíkur hf., Samherji hf., Síldarvinnslan hf., and Guðmundur Runólfsson hf. under strict confidentiality agreements. As such, the data are not publicly available and cannot be shared or accessed at any time, including upon request.

## Notes

### Competing Interest Statement

The authors have declared no competing interest.

### Summary of Updates

Add statements to clarify the main objective and assumptions of the paper Revision of statements for clarity Correct figure labels

## References

[1] Ragnar Arnason. The Icelandic fisheries : evolution and management of a fishing industry. Fishing News Books Ltd., 1 edition, 1995. doi: 10.1177/107049659500400215.

[2] Statistics Iceland. Trade in goods in the year 2023 - final data - statistics iceland, 6 2024. URL https://www.statice.is/publications/news-archive/external-trade/trade-in-goods-in-the-year-2023-final-data/#.

[3] Statistics Iceland. Export value of icelandic fisheries in 2023 - statistics iceland, 9 2024. URL https://www.statice.is/publications/news-archive/fisheries/export-value-of-icelandic-fisheries-in-2023/#.

[4] Anne Sofie Christensen, Troels Jacob Hegland, and Geir Oddsson. The icelandic itq system. Comparative Evaluations of Innovative Fisheries Management: Global Experiences and European Prospects, pages 97–118, 2009. doi: 10.1007/978-90-481-2663-7_5. URL https://link.springer.com/chapter/10.1007/978-90-481-2663-7_5.

[5] Gunnar Þórðarson, Jónas R Viðarsson, and Höfundar. Coastal fisheries in iceland skýrsla matís 12-14 coastal fisheries in iceland / smábátaveiðar við ísland. 2014. ISSN 1670-7192.

[6] Joshua K Abbott, Alan C Haynie, and Matthew N Reimer. Hidden flexibility: Institutions, incentives, and the margins of selectivity in fishing, 2015.

[7] Ricardo Alberto Cavieses Núñez, Miguel Ángel Ojeda Ruiz De La Penã, Alfredo Flores Irigollen, Manuel Rodríguez Rodríguez, and Ernesto Jardim. Deep learning models for the prediction of small-scale fisheries catches: finfish fishery in the region of “bahía magadalena-almejas”. ICES Journal of Marine Science, 75:2088–2096, 12 2018. ISSN 1054-3139. doi: 10.1093/ICESJMS/FSY065. URL https://dx.doi.org/10.1093/icesjms/fsy065.

[8] Zhongning Zhao, Ying Tian, Feng Hong, Haiguang Huang, and Shutian Zhou. Trawler fishing track interpolation using lstm for satellite-based vms traces. 2020 Global Oceans 2020: Singapore - U.S. Gulf Coast, 10 2020. doi: 10.1109/IEEECONF38699.2020.9389435.

[9] Hui Xu, Liming Song, Tianjiao Zhang, Yuwei Li, Jieran Shen, Min Zhang, and Kangdi Li. Effects of different spatial resolutions on prediction accuracy of thunnus alalunga fishing ground in waters near the cook islands based on long short-term memory (lstm) neural network model. Journal of Ocean University of China, 22: 1427–1438, 10 2023. ISSN 19935021. doi: 10.1007/S11802-023-5525-5/METRICS. URL https://link.springer.com/article/10.1007/s11802-023-5525-5.

[10] Yann LeCun, Yoshua Bengio, and Geoffrey Hinton. Deep learning. Nature, 521: 436–444, 5 2015. ISSN 14764687. doi: 10.1038/nature14539. URL https://doi.org/10.1038/nature14539.

[11] Karl Ezra Pilario, Mahmood Shafiee, Yi Cao, Liyun Lao, and Shuang Hua Yang. A review of kernel methods for feature extraction in nonlinear process monitoring. Processes, 8:1–47, 2020. ISSN 22279717. doi: 10.3390/pr8010024.

[12] Tomas Mikolov, Ilya Sutskever, Kai Chen, Greg S Corrado, and Jeff Dean. Distributed representations of words and phrases and their compositionality. In C J Burges, L Bottou, M Welling, Z Ghahramani, and K Q Weinberger, editors, Advances in Neural Information Processing Systems, volume 26. Curran Associates, Inc., 2013. URL https://proceedings.neurips.cc/paper_files/paper/2013/file/9aa42b31882ec039965f3c4923ce901b-Paper.pdf.

[13] Ashish Vaswani, Noam Shazeer, Niki Parmar, Jakob Uszkoreit, Llion Jones, Aidan N. Gomez, Łukasz Kaiser, and Illia Polosukhin. Attention is all you need. In I. Guyon, U. von Luxburg, S. Bengio, H. Wallach, R. Fergus, S. Vishwanathan, and R. Garnett, editors, Advances in Neural Information Processing Systems, volume 30. Curran Associates, Inc., 2017. URL https://proceedings.neurips.cc/paper_files/paper/2017/file/3f5ee243547dee91fbd053c1c4a845aa-Paper.pdf.

[14] Tom B Brown, Benjamin Mann, Nick Ryder, Melanie Subbiah, Jared Kaplan, Prafulla Dhariwal, Arvind Neelakantan, Pranav Shyam, Girish Sastry, Amanda Askell, Sandhini Agarwal, Ariel Herbert-Voss, Gretchen Krueger, Tom Henighan, Rewon Child, Aditya Ramesh, Daniel M Ziegler, Jeffrey Wu, Clemens Winter, Christopher Hesse, Mark Chen, Eric Sigler, Mateusz Litwin, Scott Gray, Benjamin Chess, Jack Clark, Christopher Berner, Sam McCandlish, Alec Radford, Ilya Sutskever, and Dario Amodei. Language models are few-shot learners. In Proceedings of the 34th International Conference on Neural Information Processing Systems. Curran Associates Inc., 2020. ISBN 9781713829546.

[15] Wen Song and Shigeru Fujimura. Capturing combination patterns of long- and short-term dependencies in multivariate time series forecasting. Neurocomputing, 464:72–82, 11 2021. ISSN 0925-2312. doi: 10.1016/J.NEUCOM.2021.08.100.

[16] Sepp Hochreiter and Jürgen Schmidhuber. Long short-term memory. Neural Computation, 9:1735–1780, 11 1997. ISSN 0899-7667. doi: 10.1162/neco.1997.9.8.1735. URL https://direct.mit.edu/neco/article/9/8/1735-1780/6109.

[17] Alex Graves. Long short-term memory. pages 37–45, 2012. ISSN 1860-9503. doi: 10.1007/978-3-642-24797-2_4. URL https://link.springer.com/chapter/10.1007/978-3-642-24797-2_4.

[18] Alexander Katrompas and Vangelis Metsis. Enhancing lstm models with self-attention and stateful training. Lecture Notes in Networks and Systems, 294:217–235, 2022. ISSN 23673389. doi: 10.1007/978-3-030-82193-7_14.

[19] Jian Cao, Zhi Li, and Jian Li. Financial time series forecasting model based on ceemdan and lstm. Physica A: Statistical Mechanics and its Applications, 519:127–139, 4 2019. ISSN 0378-4371. doi: 10.1016/J.PHYSA.2018.11.061.

[20] Ferdiansyah, Siti Hajar Othman, Raja Zahilah Raja Md Radzi, Deris Stiawan, Yoppy Sazaki, and Usman Ependi. A lstm-method for bitcoin price prediction: A case study yahoo finance stock market. ICECOS 2019 - 3rd International Conference on Electrical Engineering and Computer Science, Proceeding, pages 206–210, 10 2019. doi: 10.1109/ICECOS47637.2019.8984499.

[21] Adil Moghar and Mhamed Hamiche. Stock market prediction using lstm recurrent neural network. Procedia Computer Science, 170:1168–1173, 1 2020. ISSN 1877-0509. doi: 10.1016/J.PROCS.2020.03.049.

[22] Aniekan Essien and Cinzia Giannetti. A deep learning model for smart manufacturing using convolutional lstm neural network autoencoders. IEEE Transactions on Industrial Informatics, 16:6069–6078, 9 2020. ISSN 19410050. doi: 10.1109/TII.2020.2967556.

[23] Yun Bai, Jingjing Xie, Dongqiang Wang, Wanjuan Zhang, and Chuan Li. A manufacturing quality prediction model based on adaboost-lstm with rough knowledge. Computers & Industrial Engineering, 155:107227, May 2021. ISSN 0360-8352. doi: 10.1016/J.CIE.2021.107227.

[24] Changchun Liu, Haihua Zhu, Dunbing Tang, Qingwei Nie, Shipei Li, Yi Zhang, and Xuan Liu. A transfer learning cnn-lstm network-based production progress prediction approach in iiot-enabled manufacturing. International Journal of Production Research, 61:4045–4068, 6 2023. ISSN 1366588X. doi: 10.1080/00207543.2022.2056860. URL https://www.tandfonline.com/doi/abs/10.1080/00207543.2022.2056860.

[25] Md Samiul Islam, Haider Muhamed Umran, Samir M. Umran, and Mohammed Karim. Intelligent healthcare platform: Cardiovascular disease risk factors prediction using attention module based lstm. 2019 2nd International Conference on Artificial Intelligence and Big Data, ICAIBD 2019, pages 167–175, 5 2019. doi: 10.1109/ICAIBD.2019.8836998.

[26] Muhammad Awais, Mohsin Raza, Nishant Singh, Kiran Bashir, Umar Manzoor, Saif Ul Islam, and Joel J.P.C. Rodrigues. Lstm-based emotion detection using physiological signals: Iot framework for healthcare and distance learning in covid-19. IEEE Internet of Things Journal, 8:16863–16871, 12 2021. ISSN 23274662. doi: 10.1109/JIOT.2020.3044031.

[27] Dires Negash Fente and Dheeraj Kumar Singh. Weather forecasting using artificial neural network. Proceedings of the International Conference on Inventive Communication and Computational Technologies, ICICCT 2018, pages 1757–1761, 9 2018. doi: 10.1109/ICICCT.2018.8473167.

[28] Afan Galih Salman, Yaya Heryadi, Edi Abdurahman, and Wayan Suparta. Single layer & multi-layer long short-term memory (lstm) model with intermediate variables for weather forecasting. Procedia Computer Science, 135:89–98, 1 2018. ISSN 1877-0509. doi: 10.1016/J.PROCS.2018.08.153.

[29] Mohammad Safayet Hossain and Hisham Mahmood. Short-term photovoltaic power forecasting using an lstm neural network and synthetic weather forecast. IEEE Access, 8:172524–172533, 2020. ISSN 21693536. doi: 10.1109/ACCESS.2020.3024901.

[30] Zahra Karevan and Johan A.K. Suykens. Transductive lstm for time-series prediction: An application to weather forecasting. Neural Networks, 125:1–9, 5 2020. ISSN 0893-6080. doi: 10.1016/J.NEUNET.2019.12.030.

[31] Qing Li, Weidong Cai, Xiaogang Wang, Yun Zhou, David Dagan Feng, and Mei Chen. Medical image classification with convolutional neural network. 2014 13th International Conference on Control Automation Robotics and Vision, ICARCV 2014, pages 844–848, 2014. doi: 10.1109/ICARCV.2014.7064414.

[32] Khan Muhammad, Jamil Ahmad, Irfan Mehmood, Seungmin Rho, and Sung Wook Baik. Convolutional neural networks based fire detection in surveillance videos. IEEE Access, 6:18174–18183, 3 2018. ISSN 21693536. doi: 10.1109/ACCESS.2018.2812835.

[33] I. Khandokar, M. Hasan, F. Ernawan, S. Islam, and M. N. Kabir. Handwritten character recognition using convolutional neural network. Journal of Physics: Conference Series, 1918:042152, 6 2021. ISSN 1742-6596. doi: 10.1088/1742-6596/1918/4/042152. URL https://iopscience.iop.org/article/10.1088/1742-6596/1918/4/042152 https://iopscience.iop.org/article/10.1088/1742-6596/1918/4/042152/meta.

[34] Feng Yu, Qian Zhang, Jun Xiao, Yuntao Ma, Ming Wang, Rupeng Luan, Xin Liu, Yang Ping, Ying Nie, Zhenyu Tao, and Hui Zhang. Progress in the application of cnn-based image classification and recognition in whole crop growth cycles. Remote Sensing 2023, Vol. 15, Page 2988, 15:2988, 6 2023. ISSN 2072-4292. doi: 10.3390/RS15122988. URL https://www.mdpi.com/2072-4292/15/12/2988/htm https://www.mdpi.com/2072-4292/15/12/2988.

[35] Jie Hou, Badri Adhikari, and Jianlin Cheng. Deepsf: deep convolutional neural network for mapping protein sequences to folds. Bioinformatics, 34:1295–1303, 4 2018. ISSN 1367-4803. doi: 10.1093/BIOINFORMATICS/BTX780. URL https://dx.doi.org/10.1093/bioinformatics/btx780.

[36] Alok Sharma, Edwin Vans, Daichi Shigemizu, Keith A. Boroevich, and Tatsuhiko Tsunoda. Deepinsight: A methodology to transform a non-image data to an image for convolution neural network architecture. Scientific Reports 2019 9:1, 9:1–7, 8 2019. ISSN 2045-2322. doi: 10.1038/s41598-019-47765-6. URL https://www.nature.com/articles/s41598-019-47765-6.

[37] Alok Sharma, Artem Lysenko, Keith A. Boroevich, Edwin Vans, and Tatsuhiko Tsunoda. Deepfeature: feature selection in nonimage data using convolutional neural network. Briefings in Bioinformatics, 22:1–12, 11 2021. ISSN 14774054. doi: 10.1093/BIB/BBAB297. URL https://dx.doi.org/10.1093/bib/bbab297.

[38] Xingjian Shi, Zhourong Chen, Hao Wang, Dit-Yan Yeung, Wai kin Wong, and Wang chun WOO. Convolutional lstm network: A machine learning approach for precipitation nowcasting. In C Cortes, N Lawrence, D Lee, M Sugiyama, and R Garnett, editors, Advances in Neural Information Processing Systems, volume 28. Curran Associates, Inc., 2015. URL https://proceedings.neurips.cc/paper_files/paper/2015/file/07563a3fe3bbe7e3ba84431ad9d055af-Paper.pdf.

[39] Søren Kaae Sønderby, Casper Kaae Sønderby, Henrik Nielsen, and Ole Winther. Convolutional LSTM Networks for Subcellular Localization of Proteins, pages 68–80. 2015. doi: 10.1007/978-3-319-21233-3_6.

[40] Weixin Luo, Wen Liu, and Shenghua Gao. Remembering history with convolutional lstm for anomaly detection. In 2017 IEEE International Conference on Multimedia and Expo (ICME), pages 439–444, 2017. doi: 10.1109/ICME.2017.8019325.

[41] Liang Zhang, Guangming Zhu, Lin Mei, Peiyi Shen, Syed Afaq Ali Shah, and Mohammed Bennamoun. Attention in convolutional lstm for gesture recognition. In S Bengio, H Wallach, H Larochelle, K Grauman, N Cesa-Bianchi, and R Garnett, editors, Advances in Neural Information Processing Systems, volume 31. Curran Associates, Inc., 2018. URL https://proceedings.neurips.cc/paper_files/paper/2018/file/287e03db1d99e0ec2edb90d079e142f3-Paper.pdf.

[42] Ranjan Kumar Behera, Monalisa Jena, Santanu Kumar Rath, and Sanjay Misra. Colstm: Convolutional lstm model for sentiment analysis in social big data. Information Processing & Management, 58:102435, 1 2021. ISSN 03064573. doi: 10.1016/j.ipm.2020.102435.

[43] Alex Krizhevsky, Ilya Sutskever, and Geoffrey E. Hinton. Imagenet classification with deep convolutional neural networks. Communications of the ACM, 60:84–90, 5 2017. ISSN 0001-0782. doi: 10.1145/3065386.

[44] M. Mozer. A focused backpropagation algorithm for temporal pattern recognition. Complex Systems, 1989.

[45] Martin Arjovsky, Soumith Chintala, and Léon Bottou. Wasserstein gan, 2017. URL https://arxiv.org/abs/1701.07875.

[46] Zhou Wang and A.C. Bovik. Mean squared error: Love it or leave it? a new look at signal fidelity measures. IEEE Signal Processing Magazine, 26:98–117, 1 2009. ISSN 1053-5888. doi: 10.1109/MSP.2008.930649.

[47] Yukun Bao, Tao Xiong, and Zhongyi Hu. Multi-step-ahead time series prediction using multiple-output support vector regression. Neurocomputing, 129:482–493, 4 2014. ISSN 0925-2312. doi: 10.1016/J.NEUCOM.2013.09.010.

[48] Shijin Yuan, Xiaodan Luo, Bin Mu, Jing Li, and Guokun Dai. Prediction of north atlantic oscillation index with convolutional lstm based on ensemble empirical mode decomposition. Atmosphere 2019, Vol. 10, Page 252, 10:252, 5 2019. ISSN 2073-4433. doi: 10.3390/ATMOS10050252. URL https://www.mdpi.com/2073-4433/10/5/252/htm https://www.mdpi.com/2073-4433/10/5/252.

